# The bind of the burrow: space use is dominated by selection for burrow habitat over foraging habitat in an arid-adapted carnivore

**DOI:** 10.1101/2025.11.26.690549

**Authors:** Jack Thorley, Chris Duncan, Annika Herdtle, Marta Manser, Dominic Cram, Tim Clutton-Brock

## Abstract

Our understanding of habitat selection in wild vertebrates has been heavily influenced by observations of preferred foraging areas. However, foraging is only one of many ways animals interact with their environment, and preferences for habitat features that support resting, breeding, and safety, along with trade-offs between these needs, remain underexplored. These trade-offs are likely to be particularly acute in complex environments where these needs are met in different locations. Using a long-term dataset of movements and life history, we examine how preferences for foraging areas and burrow sites shape space use in Kalahari meerkats (*Suricata suricatta*). Meerkats cannot dig new sleeping and breeding burrows and must use those abandoned by other species, potentially generating trade-offs in daily space use if overnight refuges are far from optimal foraging areas. We find that space use is strongly anchored by burrow location, with burrows in calcareous pans and dry riverbed (“white sand”) habitat being preferred year-round and showing lower burrow switching rates. However, white sand areas were not preferred for foraging, and yielded lower weight gains, particularly during the dry season. As a result, meerkats faced a trade-off between optimal burrow locations and productive foraging grounds, as indicated by faster, longer, and more energetically costly travel when moving through or waking up in white sands. Our results suggest that changes in the distribution or abundance of key burrow-constructing species in desert environments may have cascading effects on the many secondary burrow-using species that depend on them for survival and reproduction. More broadly, our results highlight that limited refuge availability across the landscape can impose strong ecological constraints on animals and may restrict behavioural plasticity under environmental change, particularly if productive foraging areas shift.

## INTRODUCTION

Animals interact with their environment in diverse ways, and the habitats they select must provide the resources and conditions needed for survival and reproduction (Huey, 1991; Morris, 2003; Mayor et al., 2009; Matthiopoulos et al., 2015). Studies of habitat selection often emphasise access to food, and while foraging can clearly drive patterns of habitat use, it is only one of many functions that habitats serve. Animals also depend on refuges, breeding sites, and resting areas, among other habitat features, and their movements and habitat use will often reflect trade-offs between these competing needs (Cowlishaw, 1997; Mysterud & Ims, 1998; Godvik et al., 2009; Mayor et al., 2009; Wilson et al., 2012).

Among the non-foraging resources that animals rely on, refuges such as burrows, dens, tree cavities, or rock crevices are especially critical for many species (Hansall, 1993). They provide safety from predators (Reichman & Smith, 1987; Ebensperger & Blumstein, 2006), protection from climatic extremes (Buffenstein, 1984; Williams et al., 1999; Milling et al., 2018; Pinkert et al., 2025), and secure sites for rearing young (Wolff & Peterson, 1998; Noonan et al., 2015). Because suitable refuges are often spatially discrete, they can exert a disproportionate influence on patterns of movement, territoriality, and local population density (Macdonald & Johnson, 2015; Camp et al., 2017). For example, sleeping burrows or denning sites have been shown to constrain home ranges across mammals including carnivores, rodents, mustelids, and marsupials (Eide et al., 2001; Rosalino et al., 2005; Evans, 2008; Pomilia et al., 2015), and in primates specifically, sleeping site selection frequently shapes space use in ways that reflect strong tensions between antipredator considerations and foraging needs (Hamilton, 1982; Chapman, 1989; Markham et al., 2016).

Collectively, such findings suggest that trade-offs between preferred sheltering and feeding habitats are common. However, because most studies of habitat selection rely solely on spatial data collected by GPS-tracking devices, the behavioural drivers and energetic costs of these trade-offs are often unclear. Spatial data reveal where animals move, but they only indirectly indicate the behaviours that generate those movements, meaning that mechanisms of habitat choice (and any associated trade-offs) are typically inferred rather than demonstrated. Incorporating additional behavioural or energetic data into habitat selection studies therefore offers the potential for a richer and more ecologically grounded understanding of animals’ movement decisions (Nathan et al., 2008; Kays et al., 2015).

Secondary burrow users provide a particularly clear example of how refuge availability can impact space use and create trade-offs with access to other resources. Unlike primary burrowers, who can excavate their own sleeping and breeding burrows and may therefore be able to position themselves optimally relative to food resources (Macdonald et al., 2004), secondary burrowers depend on structures created by other species (Desmond & Savidge, 1996; Whittington-Jones et al., 2011; Di Blanco et al, 2020; Anderson et al., 2021). As a result, the availability and placement of suitable burrows are largely beyond their control and may not coincide with their foraging needs.

Here, we examine how the selection of burrow and foraging habitats influences the space use and movement patterns of meerkats (*Suricata suricatta*) in the Kalahari. Meerkats are obligate secondary burrow users, and in the Kalahari Desert, use burrows excavated by aardvarks (*Orycteropus afer*), Cape porcupines (*Hystrix africaeaustralis*), springhares (*Pedetes capensis*), and Cape ground squirrels (*Geosciurus inauris*), which they may renovate rather than digging them themselves (Figure 1). They are diurnal, desert-adapted mongooses that live in family-structured groups in which reproduction is skewed towards a dominant breeding pair (Clutton-Brock et al., 2006; Clutton-Brock & Manser, 2016). Each night, groups retreat to one of many sleeping burrows within their home range (Gall et al., 2017; Strandburg-Peshkin et al., 2020), typically using the same burrow for up to a week before switching. Burrows also serve as natal den sites during the first 3-4 weeks of pup development, when dedicated babysitters remain underground with pups while the rest of the group forages (Clutton-Brock et al., 2001). During the day, groups move cohesively throughout their home range and forage mainly on surface and shallow-dwelling invertebrates (Doolan & MacDonald, 1996; Jubber et al., 2025), occasionally making use of bolt holes and burrows in response to predators (Manser & Bell, 2004). Except under exceptional circumstances, meerkats obtain all the water they need from their food, so unlike many Kalahari mammals, their space use is not constrained by surface water (Lovegrove, 2021).

**Figure 1.**
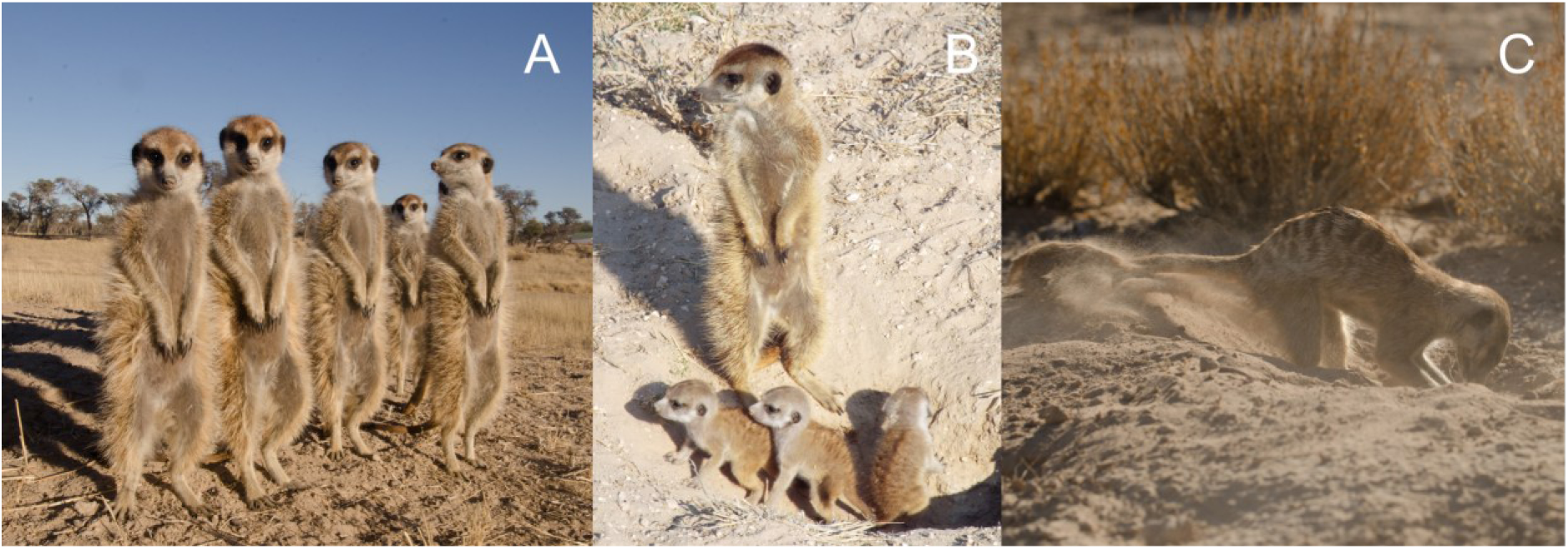
Meerkats use burrows as sleeping sites, thermal refuges, and natal dens. (A) Meerkats sunning themselves at the entrance of a sleeping burrow on a cold winter morning before departing to forage. Groups will retreat underground at night and during hot midday periods. (B) During early pup development, young remain underground under the care of babysitters while the rest of the group forages. (C) Although meerkats do not excavate their own burrows, they occasionally renovate entrances. Images (A) and (C) were taken by Kyle Finn, image (B) was taken by the first author (JT).

Our study focuses on the meerkat population at the Kuruman River Reserve in the Kalahari Desert, which has been continuously monitored for over three decades (Clutton-Brock & Manser, 2016). Because the population is habituated to close human observation, groups can be tracked at high spatial resolution (Kranstauber et al., 2020) and individuals can be weighed multiple times daily (Thorley et al., 2025a). This unique combination of data provides a rare opportunity to directly measure foraging returns and quantify the energetic consequences of space use decisions in a wild vertebrate population.

Using two decades of high-resolution movement and life-history data, we first quantify habitat selection at both landscape and local scales, distinguishing between areas used for sleeping burrow sites and those used for foraging. We then investigate the behavioural and energetic consequences of habitat choice by examining habitat-specific effects on burrow fidelity, group movement speeds, foraging track lengths, and foraging success. These complementary measures allow us to test whether meerkat groups face trade-offs between occupying habitats optimal for shelter versus those that maximise foraging returns, and to understand the mechanisms driving these decisions. By comparing the Kalahari’s wet and dry seasons, we also identify when such trade-offs are most pronounced.

## MATERIALS AND METHODS

### The habitat types

The meerkat population at the Kalahari Research Centre in South Africa (26°58’S, 21°49’E) has been continuously studied since 1993 (Clutton-Brock & Manser, 2016). The reserve is situated on the Kuruman River, which remains dry year-round except during rare, extreme rainfall events that cause flooding once every few decades. The climate features hot summers, cold winters, and significant diurnal temperature variations year-round (Thorley et al., 2025b). Annual rainfall is low (mean ± SD = 297.4 ± 135.3 mm, range = 115 to 705 mm from 1998-2023), with most precipitation falling during the months of October to April - which we refer to as the ‘wet season’.

The vegetation in and around the field site is best described as southern Kalahari duneveld (Mucina & Rutherford, 2006). Although duneveld dominates the landscape at large scales, the study area encompasses a range of finer-scale habitat types that are characteristic of the broader Kalahari region (van Rooyen et al., 2008), and that are closely related to the underlying soil (van Rooyen, 1984). Following earlier research by Harrison et al., (2021), we classified these habitats into four types based on vegetation structure and soil composition (Figure S1). For clarity, we use descriptive names throughout this study to make the classification easier for readers to follow:

1. Drie doring: calcareous white sand areas dominated by dense drie doring (*Rhigozum trichotomum*) thickets, with moderate cover of grasses.
2. White sand (pans and flats): open, sparsely vegetated flats on calcareous soils, including the riverbed and scattered pans. Vegetation is characterised by dwarf shrubs, with woody species such as *Vachellia erioloba* and *Senegalia mellifera*, as well as *Ziziphus mucronata* within the riverbed. Following rainfall, the grass layer is dominated by ‘sour grass’ *Schmidtia kalahariensis*, *Stipagrostis obtusa*, and *Chloris virgata*.
3. Red sand (low duneveld): Mixed acacia woodland on red sands, dominated by *V. erioloba*, *Vachellia haematoxylon*, and *Boscia albitrunca*, with scattered shrubs. The grass layer is well developed, including *Schmidtia kalahariensis*, *Eragrostis lehmanniana*, *Centropodia glauca*, and *Stipagrostis ciliata*, while seasonal herbs add further variation.
4. Grassland (high duneveld). Open high duneveld plains dominated by perennial grasses, particularly the dominant indicator species *Stipagrostis amabilis*. Woody plants are sparse, with occasional shrubs such as *Grewia flava* and sporadic *V. haematoxylon* trees. Various ground-spreading creepers occur, such as Gemsbok cucumber (*Acanthosicyos naudinianus*) and tsamma melon (*Citrullus lanatus*).

### The study population

Meerkat groups were visited three to four times per week throughout the project’s history, both in the morning and afternoon. Observers typically arrived at the sleeping burrow before any meerkats emerged and remained with the group throughout the morning foraging period. After emerging from the burrow each morning, meerkats typically bask for up to 90 minutes before beginning to forage. Groups forage together for 3-4 hours, after which activity declines either due to satiation or because high temperatures force them to seek shade (Habicher, 2009). In the evening, observers arrived approximately one hour before the group’s return to a burrow, which may differ from the one used the previous night.

During these observation periods, data were collected on various behavioural and life-history traits, including social and reproductive status, health status, and the expression of several cooperative behaviours (Clutton-Brock & Manser, 2016). Burrow use and group composition were also recorded. Individuals were identified using unique dye marks, and, when necessary, from passive integrated transponder (PIT) tags that were implanted in early life. On average, 13.5 ± 7.6 groups (mean ± SD; range: 6 to 21 groups) and 214.5 ± 59.4 (range: 101 to 359) known individuals were monitored per month.

Most individuals were also weighed at the start and end of each visit by coaxing them onto electronic scales in return for tiny amounts of water or hard-boiled egg. To estimate foraging returns, we used the morning weight gain, which we calculated as the difference between the pre- and post-foraging morning weights divided by the time between measurements. The average rate of morning weight gain was 6.23 ± 5.06 g/hour (mean ± SD).

During morning foraging sessions, observers also tracked group movements using handheld Garmin GPS devices. They recorded the group’s central location approximately every 15 minutes from the time they left the burrow. Between May 2002 and December 2023, 13,030 morning tracks from 61 meerkat groups were recorded, totalling 161,116 GPS fixes. From these foraging tracks we calculated the movement speed of groups, travel path lengths, and the size of their home ranges. Movement speed was calculated as the speed between successive GPS fixes (‘steps’), with groups moving at an average speed of 217.5 ± 193.8 metres per hour (mean ± SD, n = 143,922 steps). Path lengths were calculated as the summed distance of all movement steps on a given morning; the average path length was 601.1 ± 278.2 metres (n = 12710 paths). Home ranges were estimated using rolling three-month windows, including only periods in which a group had been visited at least 20 times (Thorley et al., 2025b). We calculated home ranges as the 95% autocorrelated kernel density estimate (AKDE) of all foraging locations using the `ctmm` package (Fleming & Calabrese, 2023). We retained estimates relating to four discrete quarterly periods of the year.

We conducted all our analyses in R v4.4.2 (R Core Team, 2024) using the specified packages. To ensure consistency across analyses, all datasets were restricted to the same date range (May 2002 - Dec 2023). For each statistical model, we evaluated habitat differences by estimating pairwise contrasts of the marginal means using the `emmeans` package (Lenth, 2024). We consistently compared habitat differences within each season, and for some analyses, we also compared differences across seasons for each habitat type.

### Characterising habitat selection

We used the morning foraging tracks to investigate habitat selection at both the landscape and local scale. At the landscape scale, we fitted resource selection functions (RSF), while at the local scale, we used step selection analyses (SSA) to estimate habitat selection during group movements. Both methods follow a use-availability framework, comparing locations that were used by meerkat groups to those that were available; within the home range for RSFs, and within the movement step for SSAs. By applying these complementary methods, we aimed to understand the extent to which the different habitats were selected as areas for sleeping burrows and for foraging throughout the year.

#### Landscape-scale habitat selection for sleeping burrows and foraging areas

For the RSF, we started by selecting a random point from each morning track so that each track contributed the same amount of information to the sampling procedure. These points were then used to estimate the spatial extent of each group in each season (home ranges usually expand during the wet season). To do this, we created a nonconvex boundary around all selected points for each group and season and applied a 500m buffer to account for possible movement beyond observed locations. Within each buffered extent, we randomly sampled 19 available locations per used location to represent the available habitat. Used locations were then chosen as either (1) the first GPS point of the day or (2) a randomly selected GPS point between one and four hours into foraging. This distinction allowed us to model habitat selection for burrow use versus foraging areas separately, in two models. The first location of the day is normally taken very near to the sleeping burrow (median = 13.8 metres) and therefore reflects burrow habitat selection, whereas later points represent periods of intensive foraging and inform on landscape-scale foraging preferences (Figure S2).

The two RSFs were modelled as downweighted Poisson regressions (Renner et al., 2015) using the `sdmTMB` package (Andersen et al., 2022). Habitat type was included as a fixed effect in interaction with the season, and group identity was included as a random intercept. To account for further spatial structure, we combined all spatial extents into a single mesh and incorporated the mesh as a stochastic partial differential equation (SPDE) effect (Lindgren et al., 2011). Model coefficients were interpreted as log use:availability ratios and exponentiated to estimate the relative selection intensity of the habitats (Avgar et al., 2017; Fieberg et al., 2021).

#### Local-scale habitat selection during foraging

For the step selection analysis (SSA), we selected all morning tracks lasting between one and four hours, and where consecutive GPS fixes were taken at intervals of 10 to 20 minutes (mean ± SD = 14.6 ± 1.8 minutes (mean ± SD)). Tracks lasted an average of 165.6 ± 24.8 minutes (mean ± SD). As step-selection functions require regular sampling intervals we interpolated tracks to 15-minute intervals, which represented the highest median interval in the dataset. Following interpolation, 120,011 movement steps were available for the SSA (i.e., consecutive fixes from a single track), coming from 12,866 morning tracks. For each observed movement step, we generated 19 random steps using the amt package (Signer et al., 2019), drawing from the observed distributions of step lengths (gamma distribution) and turn angles (von Mises distribution) pooled across all groups.

Used and available steps were then analysed using the Poisson regression described by Muff et al. (2020), which is equivalent to a mixed-effects conditional logistic regression model. Briefly, each observed step and its associated random steps were assigned to a unique stratum, which was fitted as a random intercept with a large, fixed variance. Habitat selection was estimated by including the habitat class at the end of the movement step as a categorical fixed effect, in addition to uncorrelated, group-specific random slopes for each habitat type to allow each group to have its own habitat-selection parameters. The models were fitted using the `glmmTMB` package (Brooks et al, 2017). We fitted separate models for the two seasons of the year. As with the RSF, model coefficients were exponentiated to reflect relative selection intensity.

### Quantifying the behavioural and energetic costs of habitat selection trade-offs

#### Habitat effects on burrow switching

If meerkat groups are selective in their choice of habitat for sleeping burrows or foraging areas, this raises questions about the mechanisms driving these preferences. Focusing first on burrow use, we hypothesised that groups would be less likely to switch burrows in the habitats that they favoured for sleeping burrows. To explore this possibility, we identified all cases where the location of a group’s sleeping burrow was known on consecutive nights, recovering 41,654 such events. Of these, 13,194 were switches (31.6%), implying that groups switch burrows once every 3 days, on average. The average distance moved between burrows was 620.8 ± 468.5 metres (mean ± SD).

We analysed the probability of burrow switching (0/1) using a generalised linear mixed effects model with binomial error distribution, implemented in the `glmmTMB` package. Fixed effects included an interaction between habitat type and the season, alongside other predictors that we expected to be important *a priori*: linear and quadratic terms for group size, a three-level categorical term for pup presence (pups being babysat at the burrow, foraging with the group, or absent), and a categorical term for breeding season. We also included an interaction between pup presence and season to allow for seasonal differences in the effect of pups. Group identity and the burrow identity were included as random effects. Pups were defined as all individuals younger than 90 days old.

#### Habitat effects on foraging success, the speed of group movements and morning track lengths

To investigate the mechanisms underlying foraging habitat selection, we analysed how habitat type influenced three complementary aspects of meerkat foraging behaviour: the rate of morning weight gain, group movement speeds and total morning track lengths. Each metric captures a different aspect of how habitat might affect foraging decisions and thus energetic outcomes. Morning weight gain, measured by the change in weight of individuals between morning and afternoon, offered a direct measure of foraging success. Movement speed served as an indicator of prey availability, with slower speeds assumed to reflect more intensive foraging effort, which is itself likely to be related to higher resource availability. Track length reflected the total distance travelled during a morning foraging session, providing information on the spatial extent of foraging effort.

All three foraging-associated metrics were analysed using linear mixed effects models and were fitted in the `nlme` package (Pinheiro & Bates, 2000; Pinheiro et al., 2024), allowing us to account for temporal autocorrelation in the data sets. The models shared a common fixed effects structure aligned with the burrow switching model, incorporating terms for the interaction between habitat type and season, linear and quadratic effects of group size, pup presence (and its interaction with season), and breeding year. We assessed the residuals for spatial autocorrelation using Moran’s I (Moran, 1950), and in each case, the value was close to zero, indicating no meaningful residual spatial structure (beyond that already captured by the model terms).

For the weight gain model, we identified the dominant habitat type encountered across the morning as the habitat variable - the habitat in which the group spent most of its time (n = 44,983 records from 1,674 individuals in 58 groups). The dominant habitat type frequently matched the habitat of the sleeping burrow used the previous night (56.7% of cases, permutation test, p < 0.001). Additional fixed effects included sex and linear and quadratic terms for individual age. We used a continuous autoregressive correlation structure of order 1 (corCAR1) to model correlation between consecutive observations within individuals, defined by the number of days since each individual’s first observation. Random effects included individual identity nested within group.

For the movement speed model (n = 143,922 steps), the habitat type at the start of each movement step served as the habitat variable. We additionally included linear and quadratic terms for time spent foraging. A first-order autoregressive correlation structure (corAR1) was applied to model autocorrelation among speeds recorded at ∼15-minute intervals. Group identity and date (nested within group) were included as random effects.

In the track length model (n = 12,710 tracks), we analysed track lengths with respect to the sleeping burrow habitat to assess whether groups starting in different habitat types travelled further. A linear variable for total tracking time was included. A continuous first-order autoregressive correlation structure (corCAR1) was included to account for temporal autocorrelation in path lengths recorded on successive days within groups. Group identity was included as a random effect.

## RESULTS

### Landscape-scale habitat selection for sleeping burrows and foraging areas

Meerkat groups displayed distinct preferences between habitats used for sleeping burrows and those used for foraging. At the population level, habitat preferences for sleeping burrows were strong and consistent throughout the year, following the order: white sand > drie doring > red sand and grassland (Figure 2C, Tables S1 & S1B). Assuming equal habitat availability, meerkat groups were 1.57 to 2.07 times more likely to select white sand over drie doring, 5.59 to 5.95 times more likely to select white sand over red sand, and 6.81 to 7.76 times more likely to select white sand over grassland, depending on the season (p < 0.001 for all pairwise contrasts).

**Figure 2.**
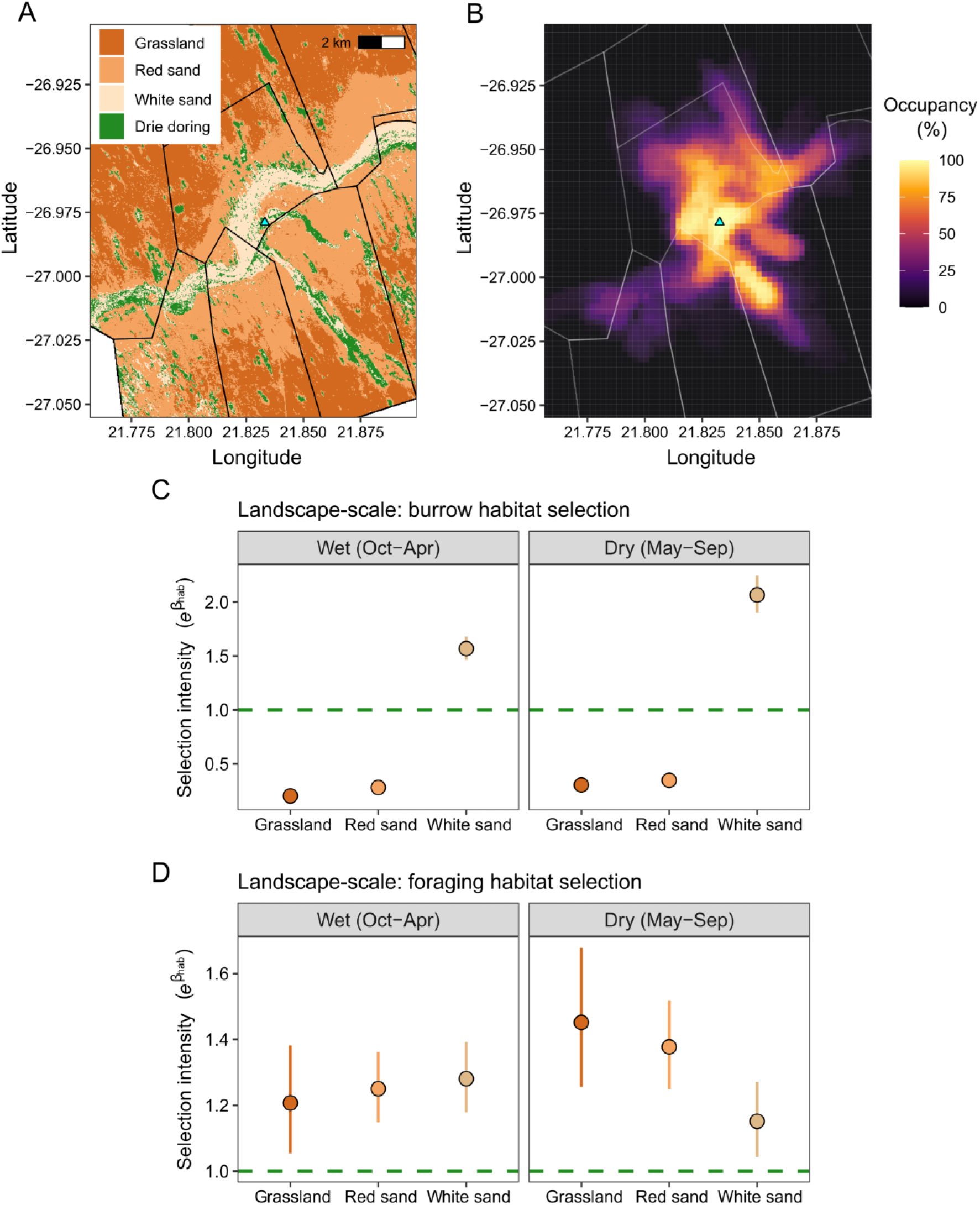
Habitat selection by meerkat groups at the Kuruman River Reserve (2002-2024). (A) The four main habitat types in and around the reserve, with the central farmhouse noted by the blue triangle. (B) The percentage use of space by established meerkat groups. Here the % refers to the percentage of periods for which each pixel overlapped the home range (95% autocorrelated kernel density estimate) of at least one meerkat group. Periods were treated as rolling 3-month windows (n = 260 periods, n = 1062 group home ranges). Habitat selection at the landscape scale, estimated through resource-selection functions, for burrows (C) and foraging areas (D). The selection intensities of the different habitat classes in wet and dry season are presented relative to the reference category of drie doring. Point estimates reflect the model-predicted means ± 95% confidence intervals.

Although white sand was the most frequently used habitat for sleeping burrows, the greater availability of red sand meant that it was used more often than drie doring, despite lower selection intensity (Table 1). Burrows in grassland areas were rarely used. The dominant role of white sand in shaping long-term burrow use - and the broader influence of sleeping burrow locations on space use - is illustrated by plotting two decades of ranging data across the study site (Figure 2B), which reveals a persistent spatial anchoring to white sand areas and limited representation of grassland in home ranges (Figure 3).

**Figure 3.**
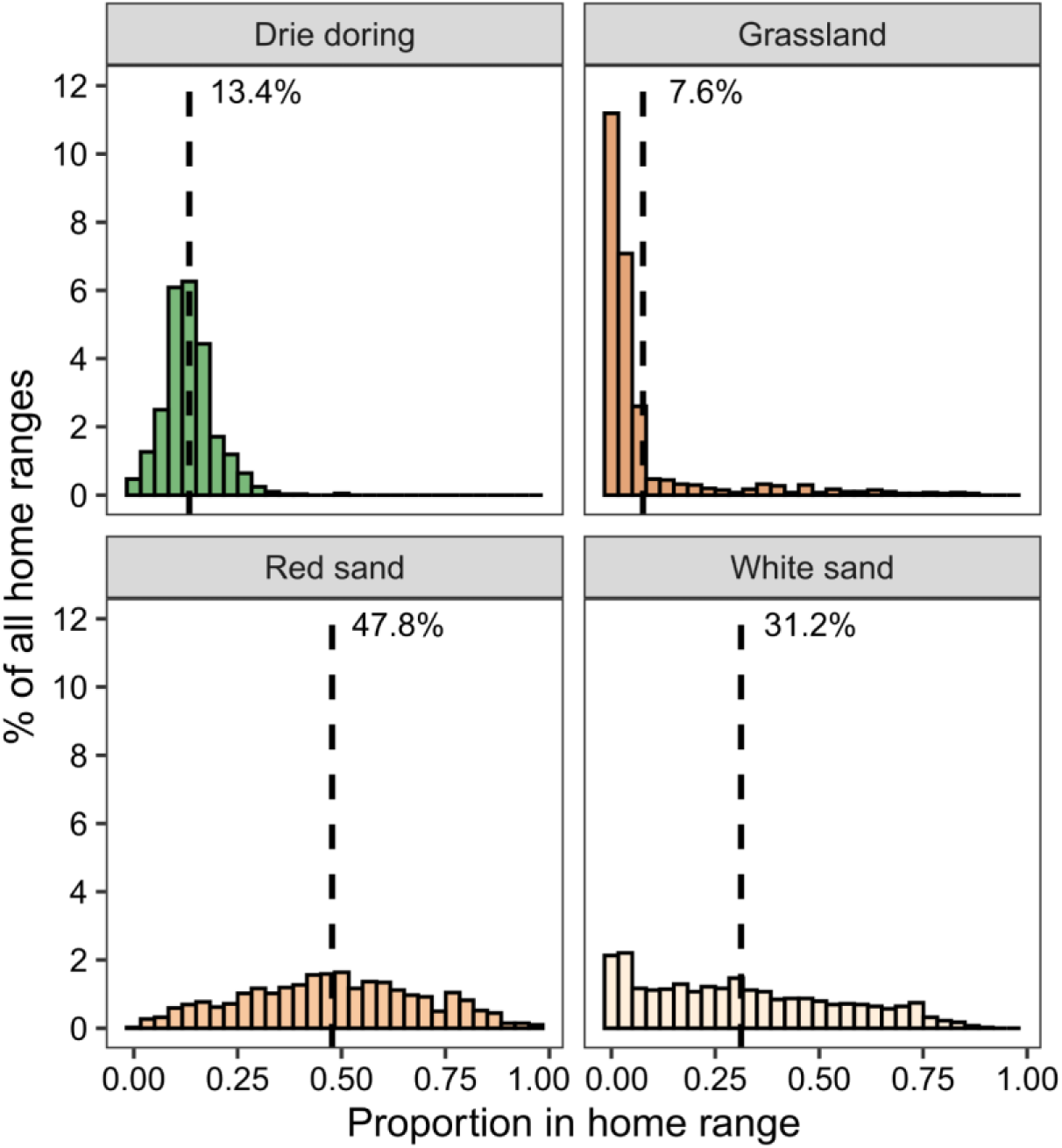
The proportion of each home range in the different habitat types. The histograms display the overall % of all home ranges, with the mean proportion in each case noted by the vertical line and the text. Home ranges were estimated as the 95% autocorrelated kernel density estimate (AKDE) of morning movement tracks in discrete 3-month windows.

**Table 1.** Estimated probability of use of the different habitats for sleeping burrows and areas of foraging in different periods of the year. Probability of use was estimated by multiplying the selection intensities from resource selection functions with the relative availability of the different habits in the landscape.

| Habitat class | Wet season<br>(Oct-Apr) | Dry season<br>(May-Sep) |
| --- | --- | --- |
| **Sleeping burrow habitat** |  |  |
| Drie doring | 16.6% | 13.2% |
| Grassland | 3.7% | 4.5% |
| Red sand | 22.1% | 21.8% |
| White sand | 57.6% | 60.5% |
| **Foraging habitat** |  |  |
| Drie doring | 9.0% | 8.5% |
| Grassland | 12.0% | 13.8% |
| Red sand | 53.5% | 55.8% |
| White sand | 25.5% | 21.8% |

Foraging habitat preferences at the landscape scale were weaker and more variable throughout the year (Figure 2D). The resource selection functions revealed one consistent pattern: drie doring was consistently avoided, with significant under-selection in all pairwise contrast (p<0.05). During the dry season (May-September), meerkats groups showed a relative preference for grassland and red sand over white sand (Tables S2 & S2B). In contrast, during the wet season (October-April), selection among grassland, red sand, and white sand was more evenly distributed, suggesting reduced habitat discrimination while foraging. Consequently, the probability of using grassland and red sand for foraging was considerably higher than for burrow placement (Table 1).

### Local-scale habitat selection during foraging

At the local scale, foraging habitat selection broadly mirrored the landscape-scale results (Figure S3, Table S3). The step selection analysis (SSA) of group movement paths indicated that drie doring was consistently avoided in both seasons. In the dry season, selection for white sand areas was also low compared to both grassland and red sand, whereas in the wet season, selection for grassland declined; white sand and red sand areas were therefore preferred at this time. The analysis also found substantial variation between groups in their foraging habitat preferences (Figure S3B).

### Habitat effects on burrow switching

The probability of meerkat groups switched burrows from one night to the next varied with both habitat type and season (Figure 4A, Table S4). Across both seasons, groups were consistently more likely to switch burrows in red sand habitats. During the dry season, the probability of burrow switching was significantly higher in red sand than in white sand (GLMM contrast ± SE = 0.19 ± 0.07, p = 0.045), with a marginally non-significant trend compared to grassland (GLMM contrast ± SE = 0.32 ± 0.13, p = 0.069). This pattern was more pronounced in the wet season, when switching in red sand was significantly higher than in drie doring (contrast = 0.26 ± 0.08, p = 0.008) and white sand (GLMM contrast ± SE = 0.24 ± 0.07, p < 0.001). In contrast, the probability of burrow switching in grassland did not differ significantly from other habitats in either season, albeit the smaller sample of grassland burrows led to wider confidence intervals.

**Figure 4.**
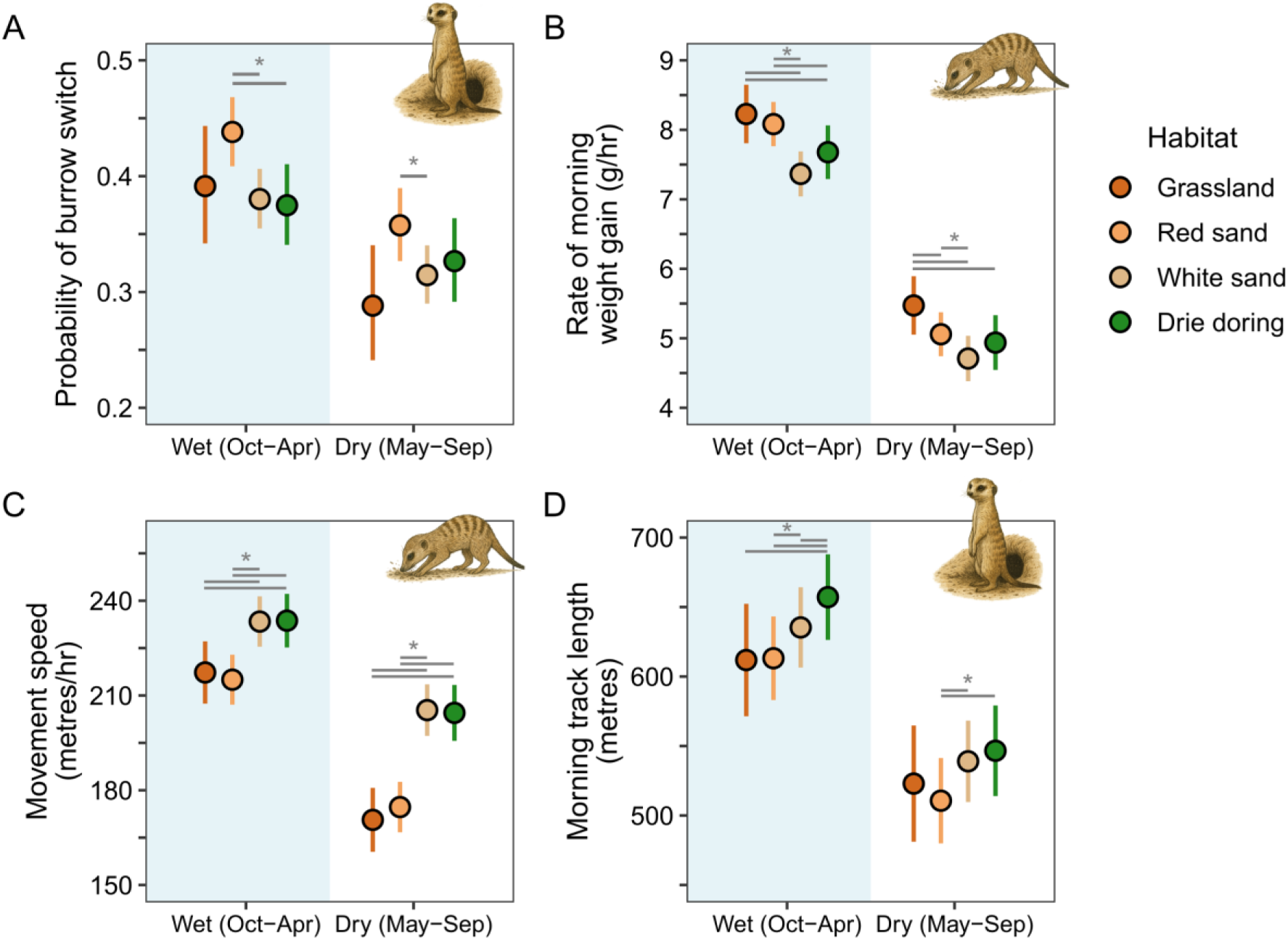
Effects of habitat type on burrow and foraging-associated metrics throughout the year. (A) Burrow switching rates, (B) morning weight gain, (C) group movement speeds, and (D) total morning track length (travel distance). Each panel displays the model-predicted means ± 95% CI for the interaction between habitat and the season of the year. Significant within-season contrasts (p < 0.05) among the habitat types are shown by the horizontal segments. The inset meerkat images indicate whether the habitat effect refers to the habitat of the previous night’s sleeping burrow or the habitat while foraging. Track length is estimated for the mean track duration of 152 minutes.

Burrow switching probability was also strongly associated with babysitting. Groups were significantly less likely to switch burrows when pups were being babysat (∼10%) compared to when pups were foraging with the group or absent (∼40%; Table S4, Figure S4B). In all cases, burrow switching was more frequent during the wet season, even though babysitting occurring more often then (34% of observations) than in the dry season (20%).

### Habitat effects on foraging success

The rate of morning weight gain varied with habitat and season, with groups generally gaining weight faster when the dominant portion of their morning was spent in grassland and red sand rather than white sand or drie doring (Figure 4B, Table S5). These differences were present in both seasons, though the magnitude and the significance of the contrasts varied. In the dry season, weight gain was higher in grassland compared to all other habitats (Table S5B, all LMM contrasts p ≤ 0.044), and in red sand versus white sand (LMM contrast ± SE = 0.35 ± 0.08 g/hour, p < 0.001). In the wet season, red sand and grassland again supported higher weight gain rates, with significant contrasts between red sand and both drie doring (LMM contrast ± SE = 0.40 ± 0.13 g/hour, p = 0.009) and white sand (LMM contrast ± SE = 0.72 ± 0.08 g/hour, p < 0.001), and between grassland and both drie doring (LMM contrast ± SE = 0.55 ± 0.19 g/hour, p = 0.023) and white sand (LMM contrast ± SE = 0.86 ± 0.17 g/hour, p < 0.001).

The rate of morning weight gain also varied with individual- and group-level factors (Figure S5, Table S5). Females gained weight faster than males (LMM estimate ± SE = 0.57 ± 0.07 g/hour, t-value 8.07, p < 0.001), and weight gain followed a non-linear age pattern, increasing with age before declining in older individuals. Individuals also gained more weight when pups were being babysat, while group size had little effect.

### Habitat effects on the speed of group movements and morning track lengths

Across both seasons, groups moved fastest through white sand and drie doring, and slowest through grassland and red sand (Figure 4C, Table S6 & S6B). These differences were most pronounced during the dry season, when groups moved between 29.8 and 34.7 metres/hour faster in white sand and drie doring compared to red sand or grassland, corresponding to a 14.1 to 16.8% increase in speed (p < 0.001 in all LMM contrasts). In the wet season, when movement speeds were generally higher, the contrasts between the habitats were smaller but still significant. Groups also moved significantly faster shortly after leaving the burrow in the morning, when dependent pups were being babysat, and in larger groups (Table S6, Figure S6).

Meerkat groups also covered longer distances over the course of the morning when emerging from sleeping burrows in white sand and drie doring, though effect sizes were sometimes small and not always statistically significant (Figure 4D, Table S7 & S7B). This pattern was strongest in the wet season, when track lengths from drie doring burrows were longer than those from red sand (LMM contrast ± SE = 43.3 ± 9.2 metres, p < 0.001), grassland (LMM contrast ± SE = 44.1 ± 16.8 metres, p = 0.044), and white sand (LMM contrast ± SE = 22.6 ± 7.9 metres, p = 0.035), while those from white sand burrows also exceeded those from red sand burrows (contrast = 20.7 ± 7.7 metres, p = 0.035).

A similar, though less marked, pattern occurred in the dry season, with groups emerging from red sand burrows covering shorter distances than those from drie doring (LMM contrast ± SE = −35.9 ± 11.1 metres, p = 0.007) or white sand (LMM contrast ± SE = −26.9 ± 8.7 metres, p = 0.010). Track lengths were positively associated with group size (Table S7, Figure S7A) and were reduced when dependent pups were present, and especially when pups were foraging with the group (Table S7, Figure S7B). Morning tracks were longer during the wet season, in line with the higher movement speeds at this time.

## DISCUSSION

Our study shows that meerkat space use in the Kalahari is shaped more strongly by the distribution of suitable burrow habitat than by foraging opportunities. Using two decades of fine-scale tracking and life-history data, we found that groups consistently anchored their ranges to calcareous white sand habitats, which provided the preferred locations for sleeping burrows, despite these areas offering lower foraging returns than alternative habitats. Foraging habitat selection was more flexible and responsive to seasonal conditions but remained secondary to the spatial constraints imposed by burrows. This trade-off was evident in faster and longer tracks in white sands and drie doring habitats, reflecting their lower foraging value. Our results indicate that decisions about where to sleep and rear offspring play a primary, if not the main, role in shaping the movement and spatial behaviour of this arid-adapted carnivore. Given that many other carnivores also rely on burrows or dens they do not construct themselves (Noonan et al., 2015), such trade-offs are likely a common feature of carnivore ecology.

### Sleeping burrow habitat dominates space use

Previous work on our population has described aspects of burrow dynamics (Dyble et al., 2019; Strandburg-Peshkin et al., 2020) and foraging habitat selection (Turbé, 2006; Bateman et al., 2015), but ours is the first study to integrate these dimensions to examine their joint influence on space use. Across seasons and years, meerkats consistently preferred burrows in white sand areas, showed lower switching probabilities there, and maintained long-term associations with these patches. The reasons for higher burrow fidelity in white sand areas - including calcareous pans, flats, and the dry riverbed - remain uncertain. One possibility is that soil properties in these areas allow primary excavators to construct deeper, more stable burrows with multiple entrances and larger chamber systems, providing secondary users like meerkats improved thermal buffering and greater space; the latter may be particularly important given that the average meerkat group numbers around 14 individuals (Thorley et al., 2025a). Alternatively, higher burrow fidelity may reflect the spatial clustering of burrows rather than structural differences related to soil characteristics. Whatever the cause, the strong association of meerkats with white sand habitats underscores the disproportionate importance of sleeping burrows in structuring their space use.

Strong reliance on refuges is likely a common feature of many animals in arid environments, where alternative shelters are scarce and extreme temperatures promote belowground activity across numerous taxa (Kinlaw, 1999; Oliveira et al., 2024; Pinkert et al., 2025). Supporting this view, refuge availability strongly shapes movement in other arid-adapted species: the Australian plains mouse (*Pseudomys australis*) confines its dry-season home range to cracks within specific clay habitats (Young et al., 2017), degus exhibit more bursty locomotion in open areas (Vásquez et al., 2002), and desert-living baboons prioritise low-risk food patches near cliffs and trees, even when these offer poorer foraging returns (Cowlishaw, 1997). These studies underscore how refuge placement governs daily movement flexibility and home-range structure. Our work integrates behavioural, physiological, and spatial data to provide a mechanistic perspective on space use. While such extensive datasets are uncommon, in our case relying on habituated animals and extensive human effort, the advent of multi-sensor biologgers now allows integrated behavioural and spatial analyses. Their application of such methods will likely be especially valuable for understanding habitat selection in arid systems as warming climates intensify refuge-related constraints.

### Foraging space use is more flexible and secondary

Multiple lines of evidence indicate that foraging habitat selection was weaker and more seasonally dependent than burrow site selection. Groups preferentially foraged in red sand and grassland during the dry season, a preference that diminished in the wet season when all habitat types offered higher foraging returns. Red sand and grassland areas were also associated with reduced group movement speeds and higher morning body weight gains, suggesting greater prey availability. Yet, despite these energetic advantages, groups rarely used burrows in the most productive foraging habitats and instead often travelled from preferred refuges to reach them.

While food availability probably drives the habitat-specific foraging decisions of meerkats, vegetation structure and predation risk may also play a role. The white sand areas at our field site are extremely open, especially in the dry season (Figure S1), which may increase exposure to aerial or terrestrial predators and render them less attractive foraging spots. Indeed, like many small mammals, meerkats prefer to remain close to vegetative cover while foraging (Le Roux et al., 2009). The dry riverbed also hosts most of the tall trees in the area, which provide potential perching sites for the Kalahari’s many raptors, and possibly increasing vigilance demands. Conversely, the avoidance of drie doring habitats may reflect the low permeability of areas that are densely covered by this shrub. Other landscape features, such as bolt hole density, may further influence foraging decisions, and the presence of neighbouring groups likely plays an additional role (Bateman et al., 2015; Kranstauber et al., 2020). Separating the relative contributions of food distribution, landscape permeability, and predation risk, could be an important avenue for future research.

### Energetic trade-offs and life-history consequences

Our results support the idea that trade-offs between refuges and food acquisition can be pronounced in obligate secondary burrow users, which cannot relocate or modify refuges to align with food availability. Accordingly, groups with burrows in white sand or drie doring habitats travelled further during morning foraging bouts, and those that spent most of their time foraging in these habitats showed lower weight gains. Such reductions in daily weight gain are expected to cascade into effects on reproductive success and survival (Thorley et al., 2025a), though the relatively small scale of white sand patches seems to allow most established groups to balance these needs effectively most of the time.

While not explicitly tested here, we expect that the foraging-shelter trade-off becomes more difficult to manage during pup rearing, when fidelity to sleeping burrows peaks due to their role as denning sites. Groups then effectively become central-place foragers, with reduced flexibility to adjust their ranging in response to changing foraging conditions. Our long-term data further suggest that the trade-off may be intensifying, as morning foraging returns have declined over the past two decades (Figure S5D), mirroring concurrent declines in adult body mass (Thorley et al., 2025b). Together, these patterns highlight the growing influence of climate warming and more frequent droughts on foraging success, trends that if they continue, will amplify the constraint imposed by the spatial separation of refuges and foraging habitats.

### Broader spatial and ecological implications

The spatial dynamics we uncover are likely to scale beyond group ranges to also impact the connectivity of meerkat populations across the wider landscape. Much of the species’ distribution lies within the arid savannah biome of the Kalahari, where duneveld predominates (referred to in our study as grassland). In contrast, the white sand pans and dry riverbeds that provide their favoured burrow habitat occupy a much smaller fraction of the landscape (Thomas & Shaw, 1991). It is therefore unsurprising on reflection that the two major ecological studies of meerkats have been situated on the dry riverbeds of the Nossob River in the Kgalagadi Transfrontier Park, and the Kuruman River; sites which provide abundant sleeping burrow habitat. Though speculative, our results suggest that large portions of the Kalahari may present suboptimal habitat for meerkats, with the patchy distribution of sleeping burrow habitat constraining dispersal opportunities and potentially limiting gene flow. Population genetic studies could test this idea.

Beyond the Kalahari, meerkats occur throughout the Nama-Karoo biome of South Africa. Whether these populations experience similar trade-offs between burrow availability and foraging opportunities remains unknown, as even basic ecological descriptions are lacking. Comparisons across populations, including those in truly arid regions such as the Namib Desert, would clarify how landscape structure, sleeping burrow habitat, and foraging preferences determine space use across the species’ range.

Finally, our study highlights that shifts in the abundance or distribution of primary burrowers could disproportionately affect secondary burrow users, reinforcing the role of primary excavators as keystone species. While existing burrows may persist for some time, they are not permanent, and changes in their availability may cascade through ecosystems. In the Kalahari, this is particularly concerning given recent evidence that climate change is already impacting the region’s most well-known ecosystem engineer, the aardvark (Rey et al., 2017).

## Conclusion

In summary, our study demonstrates that in a desert-adapted, secondary burrow-using carnivore, the spatial distribution of refuges outweighs that of food in shaping space use, movement, and energetic outcomes. Burrows anchor home ranges and impose trade-offs that reveal the often-underappreciated role of non-foraging habitat features in driving animal behaviour. Our findings underscore the need for habitat selection frameworks that explicitly integrate multiple resource types and consider the constraints they impose on behavioural flexibility. As environmental change reshapes the distribution of both refuges and foraging resources, understanding how species navigate these trade-offs will be essential for predicting their persistence and distribution in arid ecosystems.

## ACKNOWLEDGEMENTS

We are grateful to the Kalahari Research Trust and the Kalahari Meerkat Project for providing access to the facilities and habituated animals at the Kuruman River Reserve, and to the neighbouring landowners for permitting access to their properties. We owe sincere thanks to the many managers, volunteers, students, and researchers whose efforts made the long-term data collection and curation possible. We extend special thanks to Walter Jubber and Tim Vink for their dedication in managing the reserve and its databases. We also acknowledge the logistical support from the Mammal Research Institute and the ethics committee at the University of Pretoria, as well as the Northern Cape Department of Environment and Nature Conservation for granting research permission (FAUNA 0930/2022).

## AUTHOR CONTRIBUTIONS

*Conceptualisation:* JT, supported by CD, DC and TC-B. *Data curation*: TC-B and MM. *Formal analysis*: JT, supported by CD and AH. *Funding acquisition:* TC-B and MM. *Investigation*: All authors. *Project Administration*: TC-B. *Writing*: JT. *Review and editing*: All authors.

## OPEN RESEARCH STATEMENT

Data and reproducible R code are available at https://github.com/JThor1990/Meerkat-habitat-selection. The track data are too large for Github and contain a large historic data set. Reasonable requests for these data will be considered by the Kuruman Research Centre’s principal investigators.

## ANIMAL ETHICS STATEMENT

The long-term data used in this study were collected under permits granted by the ethics committee of the University of Pretoria, South Africa (EC047-16, EC010-13, SOP029-12), and adhered to the standards outlined in the ASAB/ABS guidelines for the Treatment of Animals in Behavioural Research and Training (ASAB, 2012).

