## Supplementary Material for "The bind of the burrow: space use is dominated by selection for burrow habitat over foraging habitat in an arid-adapted carnivore"

#### *Kalahari habitats*

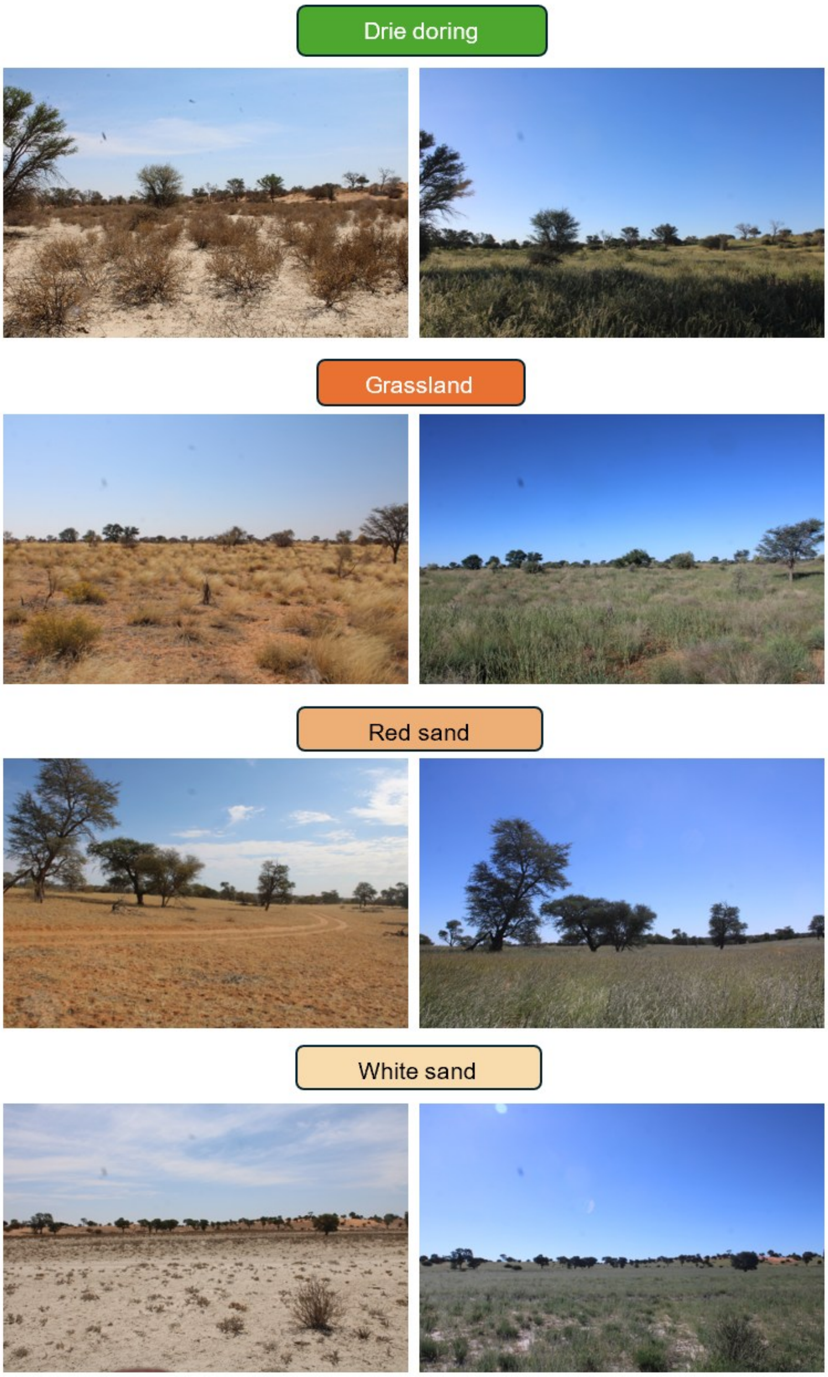


**Figure S1. Photographs of the four main habitat types encountered by meerkat groups at our study site. In each case, we provide a photo at the end of the dry season, and a location-matched photo taken at the end of typical wet season.**

#### *Occupancy of different habitat types across the morning foraging period*

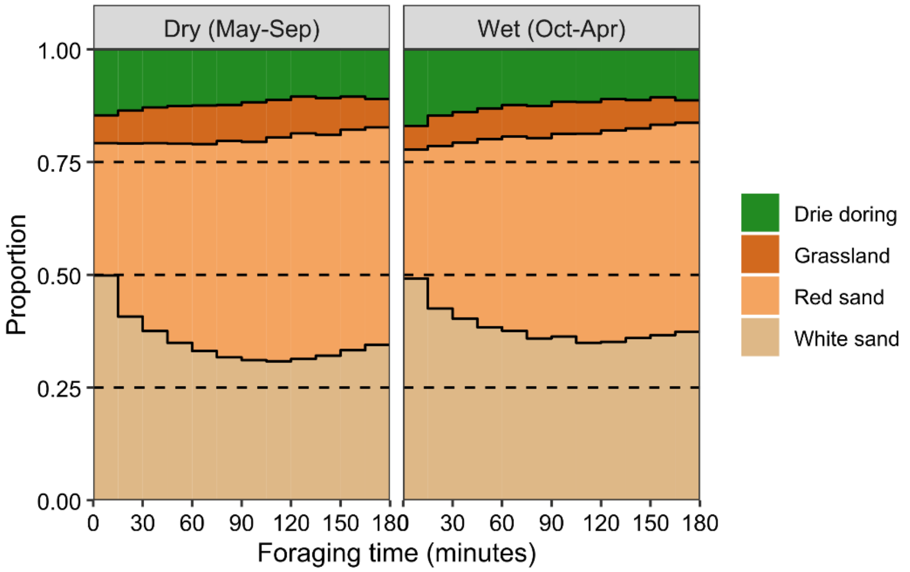


**Figure S2. The proportion of time spent in different habitats across the morning's foraging window (~ 3 hours) in the wet (Oct-Apr) and the dry season (May-Sep). Foraging commences when groups leave the immediate vicinity of their sleeping burrow. The fact that relative use of drie doring and white sand areas is higher first thing in the morning supports resource selection and step selection analyses showing that burrow use is concentrated in these habitat types throughout the year.**

### *Landscape-scale habitat selection via resource selection functions (RSF)*

**Table S1. Model output table for the resource selection analysis of sleeping burrow habitat. RSF modelled as a downweighted Poisson regression with a random intercept of group identity (std dev = 0.42) and spatial random field fitted as a SPDE (std dev = 2.96). Model estimates are provided on the [log] link scale. The pairwise contrasts for period-specific habitat differences are presented in Table S1B. The intercept represents the reference level of 'drie doring' in the dry season period of 'May-Sep'. The wet season is thus 'Oct-Apr'.**

| Term | Estimate [Std Error] |
|---|---|
| Intercept | 2.49 [0.27] |
| Habitat (Grassland) | -1.19 [0.09] |
| Habitat (Red sand) | -1.06 [0.05] |
| Habitat (White sand) | 0.73 [0.04] |
| Season (Wet) | 0.23 [0.05] |
| Habitat (Grassland) : Season (Wet) | -0.41 [0.09] |
| Habitat (Red sand) : Season (Wet) | -0.21 [0.06] |
| Habitat (White sand) : Season (Wet) | -0.28 [0.05] |

**Table S1B. Pairwise habitat contrasts from the model presented in Table S1. For each season, the lower triangle of the matrix corresponds to the estimated difference in the marginal means [std error] of the habitat types (row versus column), and the upper triangle indicates the associated p-value. p values ≤ 0.05 are indicated in bold.**

| **Aug-Oct (DRY)** | Drie doring | Grassland | Red sand | White sand |
|---|---|---|---|---|
| Drie doring | -- | **<0.001** | **<0.001** | **<0.001** |
| Grassland | -1.19 [0.09] | -- | 0.33 | **<0.001** |
| Red sand | -1.06 [0.05] | 0.13 [0.08] | -- | **<0.001** |
| White sand | 0.73 [0.04] | 1.92 [0.08] | 1.78 [0.04] | -- |
| **Nov-Jan (WET)** | Drie doring | Grassland | Red sand | White sand |
| Drie doring | -- | **<0.001** | **<0.001** | **<0.001** |
| Grassland | -1.60 [0.08] | -- | **<0.001** | **<0.001** |
| Red sand | -1.27 [0.04] | 0.32 [0.07] | -- | **<0.001** |
| White sand | 0.45 [0.03] | 2.05 [0.08] | 1.72 [0.04] | -- |

**Table S2. Model output table for the resource selection analysis of foraging habitat. RSF modelled as a downweighted Poisson regression with a random intercept of group identity (std dev = 0.36) and spatial random field fitted as a SPDE (std dev = 2.47). Model estimates are provided on the [log] link scale. The pairwise contrasts for period-specific habitat differences are presented in Table S2B. The intercept represents the reference level of 'drie doring' in the dry season period of 'May-Sep'. The wet season is thus 'Oct-Apr'.**

| Term | Estimate [Std Error] |
|---|---|
| Intercept | 1.27 [0.72] |
| Habitat (Grassland) | 0.37 [0.07] |
| Habitat (Red sand) | 0.32 [0.05] |
| Habitat (White sand) | 0.14 [0.05] |
| Season (Wet) | 0.02 [0.05] |
| Habitat (Grassland) : Season (Wet) | -0.18 [0.08] |
| Habitat (Red sand) : Season (Wet) | -0.10 [0.06] |
| Habitat (White sand) : Season (Wet) | 0.11 [0.06] |

**Table S2B. Pairwise habitat contrasts from the model presented in Table S2. For each season, the lower triangle of the matrix corresponds to the estimated difference in the marginal means [std error] of the habitat types (row versus column), and the upper triangle indicates the associated p-value. p values ≤ 0.05 are indicated in bold.**

| **May-Sep (DRY)** | Drie doring | Grassland | Red sand | White sand |
|---|---|---|---|---|
| Drie doring | -- | **<0.001** | **<0.001** | **0.025** |
| Grassland | 0.37 [0.07] | -- | 0.82 | **0.004** |
| Red sand | 0.32 [0.05] | -0.05 [0.06] | -- | **<0.001** |
| White sand | 0.14 [0.05] | -0.23 [0.07] | -0.18 [0.04] | -- |

| **Oct-Apr (WET)** | Drie doring | Grassland | Red sand | White sand |
|---|---|---|---|---|
| Drie doring | -- | **0.032** | **<0.001** | **<0.001** |
| Grassland | 0.19 [0.07] | -- | 0.93 | 0.80 |
| Red sand | 0.22 [0.04] | 0.04 [0.07] | -- | 0.92 |
| White sand | 0.25 [0.04] | 0.06 [0.07] | 0.02 [0.04] | -- |

### *Local-scale habitat selection via step selection analysis (SSA)*

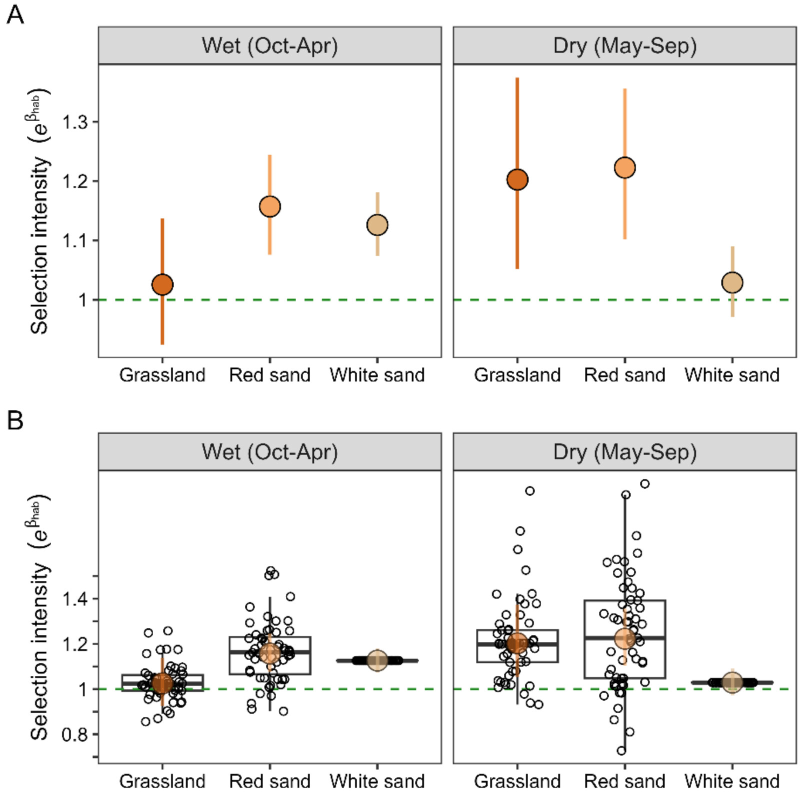


**Figure S3. Local-scale habitat selection of meerkats while foraging. (A) The selection intensity of the different habitat classes in each period is presented relative to the reference category of drie doring. Point estimates reflect the mean ± 95% confidence intervals at the population-level. The dashed horizontal line indicates no preference relative to drie doring. (B) Group-specific variation in habitat selection. The boxplots and overlaid points display individual group estimates, while the larger points display the population-level effect (mean ± 95%). The dashed horizontal line indicates no preference relative to drie doring in either season.**

**Table S3. Pairwise habitat contrasts from the step selection analyses presented for the SSA in Figure S4. Each season was modelled separately. For each season, the lower triangle of the matrix corresponds to the estimated difference in the marginal means [std error] of the habitat types (row versus column), and the upper triangle indicates the associated p-value. p values ≤ 0.05 are indicated in bold.**

| **May-Sep (DRY)** | Drie doring | Grassland | Red sand | White sand |
|---|---|---|---|---|
| Drie doring | -- | **0.034** | **<0.001** | 0.77 |
| Grassland | 0.18 [0.07] | -- | 1.00 | 0.077 |
| Red sand | 0.20 [0.05] | 0.02 [0.07] | -- | **0.002** |
| White sand | 0.03 [0.03] | -0.16 [0.07] | -0.17 [0.05] | -- |
| **Oct-Apr (WET)** | Drie doring | Grassland | Red sand | White sand |
| Drie doring | -- | 0.96 | **<0.001** | **<0.001** |
| Grassland | 0.03 [0.05] | -- | 0.094 | 0.24 |
| Red sand | 0.15 [0.04] | 0.12 [0.05] | -- | 0.84 |
| White sand | 0.12 [0.02] | 0.09 [0.05] | -0.03 [0.03] | -- |

### *Burrow switching*

**Table S4. Model output for the Generalised linear mixed effects model of burrow switching rates. The response was modelled to a binomial error distribution with a random intercept of group (variance = 0.04, std dev = 0.20) and burrow identity (variance = 0.32, std dev = 0.56). Model estimates are provided on the [logit] link scale. The pairwise contrasts for season-specific habitat differences are presented in Table S4B. The intercept represents the reference habitat level of 'drie doring' in the dry season 'May-Sep', when a group was babysitting pups. We modelled the year ('breeding season') as a fixed rather than a random effect so that we could estimate year-specific means directly (Figure S4C). Continuous variables were standardised and centred for model fitting. p values ≤ 0.05 are indicated in bold.**

| Term | Estimate [Std Error] | z-value | p-value |
|---|---|---|---|
| Intercept | -2.59 [0.17] | - | - |
| Habitat (Grassland) | -0.18 [0.14] | -1.29 | 0.20 |
| Habitat (Red sand) | 0.14 [0.09] | 1.47 | 0.14 |
| Habitat (White sand) | -0.06 [0.09] | -0.65 | 0.52 |
| Season (Wet) | 0.43 [0.09] | 4.71 | **<0.001** |
| Habitat (Grassland) : Season (Wet) | 0.25 [0.14] | 1.86 | 0.063 |
| Habitat (Red sand) : Season (Wet) | 0.13 [0.08] | 1.62 | 0.11 |
| Habitat (White sand) : Season (Wet) | 0.08 [0.06] | 1.20 | 0.23 |
| Group size | 0.03 [0.02] | 1.41 | 0.16 |
| Group size$^2$ | -0.01 [0.01] | -1.36 | 0.17 |
| Pup presence (neither) | 1.97 [0.07] | 27.68 | **<0.001** |
| Pup presence (foraging) | 1.96 [0.08] | 25.69 | **<0.001** |
| Pup presence (neither) : Season (Wet) | -0.31 [0.08] | -3.72 | **<0.001** |
| Pup presence (foraging) : Season (Wet) | -0.30 [0.09] | -3.45 | **<0.001** |
| Year (2002/2003) | 0.41 [0.15] | 2.80 | **0.005** |
| Year (2003/2004) | 0.39 [0.15] | 2.62 | **0.009** |
| Year (2004/2005) | 0.35 [0.15] | 2.31 | **0.021** |
| Year (2005/2006) | 0.29 [0.15] | 1.88 | 0.060. |
| Year (2006/2007) | 0.29 [0.15] | 1.96 | **0.050** |
| Year (2007/2008) | 0.45 [0.15] | 2.99 | **0.003** |
| Year (2008/2009) | 0.2 [0.15] | 1.38 | 0.17 |
| Year (2009/2010) | 0.43 [0.15] | 2.91 | **0.004** |
| Year (2010/2011) | 0.58 [0.15] | 3.92 | **<0.001** |
| Year (2011/2012) | 0.25 [0.15] | 1.72 | 0.085 |
| Year (2012/2013) | 0.36 [0.15] | 2.44 | **0.015** |
| Year (2013/2014) | 0.47 [0.15] | 3.14 | **0.002** |
| Year (2014/2015) | 0.62 [0.15] | 4.14 | **<0.001** |
| Year (2015/2016) | 0.90 [0.15] | 5.89 | **<0.001** |
| Year (2016/2017) | 0.61 [0.15] | 4.06 | **<0.001** |
| Year (2017/2018) | 0.53 [0.15] | 3.52 | **<0.001** |
| Year (2018/2019) | 0.74 [0.15] | 4.92 | **<0.001** |
| Year (2019/2020) | 0.65 [0.15] | 4.24 | **<0.001** |
| Year (2020/2021) | 0.33 [0.16] | 2.09 | **0.037** |
| Year (2021/2022) | 0.45 [0.16] | 2.86 | **0.004** |
| Year (2022/2023) | 0.34 [0.16] | 2.20 | **0.028** |
| Year (2023/2024) | 0.37 [0.18] | 2.10 | **0.036** |

**Table S4B. Pairwise habitat contrasts from the burrow switching model presented in Table S4. For each period, the lower triangle of the matrix corresponds to the estimated difference in the marginal means [std error] and the upper triangle indicates the associated p-value. p values ≤ 0.05 are indicated in bold. Note that the contrasts are here on the [logit] link scale.**

| **May-Sep (DRY)** | Drie doring | Grassland | Red sand | White sand |
|---|---|---|---|---|
| Drie doring | -- | 0.57 | 0.46 | 0.92 |
| Grassland | -0.18 [0.14] | -- | 0.069 | 0.76 |
| Red sand | 0.14 [0.09] | 0.32 [0.13] | -- | **0.045** |
| White sand | -0.06 [0.09] | 0.13 [0.13] | -0.19 [0.07] | -- |

| **Oct-Apr (WET)** | Drie doring | Grassland | Red sand | White sand |
|---|---|---|---|---|
| Drie doring | -- | 0.94 | **0.008** | 0.99 |
| Grassland | 0.07 [0.12] | -- | 0.32 | 0.98 |
| Red sand | 0.26 [0.078] | 0.19 [0.11] | -- | **<0.001** |
| White sand | 0.02 [0.08] | -0.05 [0.11] | -0.24 [0.07] | -- |

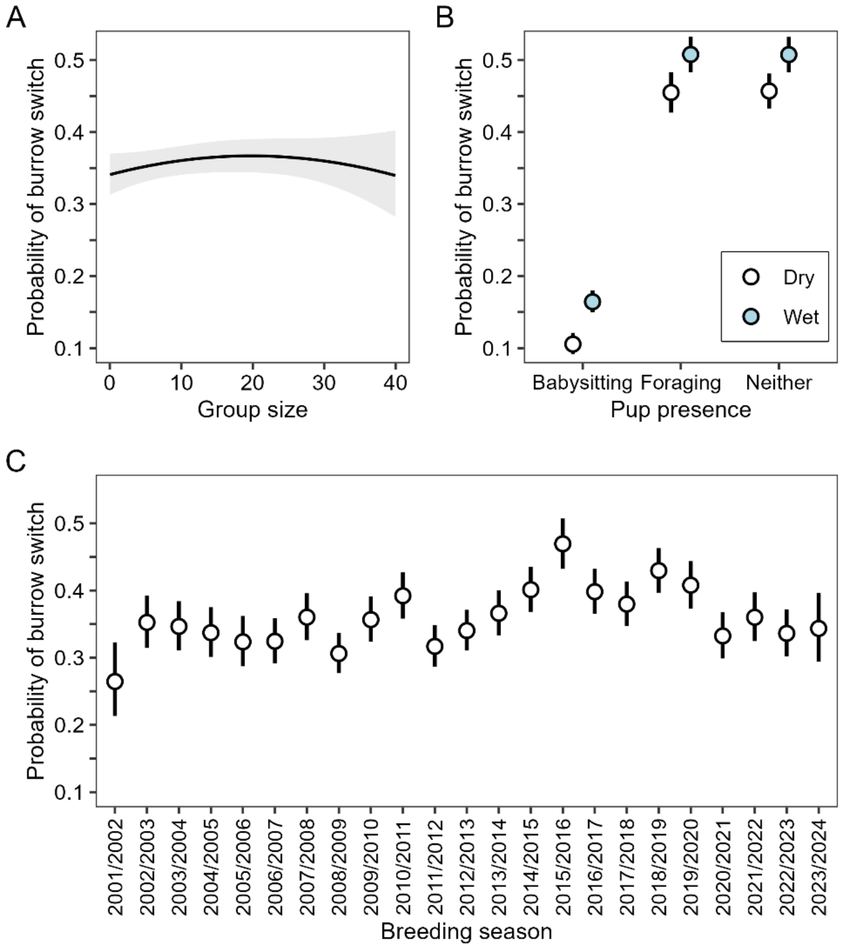


**Figure S4. Variation in the probability of burrow switching according to (A) group size, (B) the presence of pups at the group, and (C) the breeding season (C), as predicted by a generalised linear mixed effects model with a binomial error distribution. The presence of pups was either due to their permanent presence at burrow in early life, when they are babysat by one or more adult helpers, or later, once they begin foraging with the group. All panels display the predicted mean ± 95% confidence intervals.**

### *Rate of morning weight gain*

**Table S5. Model output for the linear mixed effects model of the average rate of morning weight gain. The pairwise contrasts for period-specific habitat differences are presented in Table S6B. The intercept represents the reference habitat cluster of 'drie doring' in the dry season period of 'May-Sep', for adult females. We modelled the year as a fixed rather than a random effect so that we could estimate year-specific means directly (Figure S5D). The model included a random intercept of individual identity (std dev = 0.96) nested within group identity (std dev = 1.03), and an autocorrelation parameter (continuous autoregressive correlation of order 1, corCAR1) was applied to the repeated observations individuals within their groups over time (phi = 0.20). Continuous variables were standardised and centred for model fitting. Residual std dev = 4.59. p values ≤ 0.05 are indicated in bold.**

| Term | Estimate [Std Error] | t-value | p-value |
|---|---|---|---|
| Intercept | 6.58 [0.38] | - | - |
| Habitat (Grassland) | 0.53 [0.20] | 2.66 | **0.008** |
| Habitat (Red sand) | 0.12 [0.14] | 0.88 | 0.38 |
| Habitat (White sand) | -0.23 [0.14] | -1.61 | 0.107 |
| Season (Wet) | 3.06 [0.20] | 15.07 | **<0.001** |
| Habitat (Grassland) : Season (Wet) | 0.02 [0.26] | 0.06 | 0.95 |
| Habitat (Red sand) : Season (Wet) | 0.28 [0.18] | 1.54 | 0.12 |
| Habitat (White sand) : Season (Wet) | -0.08 [0.19] | -0.44 | 0.66 |
| Group size | 0.08 [0.05] | 1.65 | 0.098 |
| Group size$^2$ | -0.00 [0.03] | -0.06 | 0.95 |
| Age | 0.48 [0.05] | 10.12 | **<0.001** |
| Age$^2$ | -0.15 [0.02] | -8.38 | **<0.001** |
| Sex (Male) | -0.57 [0.07] | -8.07 | **<0.001** |
| Pup presence (neither) | -0.74 [0.10] | -7.24 | **<0.001** |
| Pup presence (foraging) | -0.76 [0.15] | -5.21 | **<0.001** |
| Pup presence (neither) : Season (Wet) | -0.34 [0.13] | -2.69 | **0.007** |
| Pup presence (foraging) : Season (Wet) | -0.68 [0.17] | -3.87 | **<0.001** |
| Year (2002/2003) | 0.87 [0.31] | 2.80 | **0.005** |
| Year (2003/2004) | 0.35 [0.32] | 1.08 | 0.28 |
| Year (2004/2005) | -0.20 [0.33] | -0.62 | 0.54 |
| Year (2005/2006) | -0.71 [0.35] | -2.04 | **0.041** |
| Year (2006/2007) | 0.95 [0.33] | 2.91 | **0.004** |
| Year (2007/2008) | -1.27 [0.35] | -3.62 | **<0.001** |
| Year (2008/2009) | -0.19 [0.35] | -0.53 | 0.60 |
| Year (2009/2010) | -0.78 [0.35] | -2.22 | **0.026** |
| Year (2010/2011) | -1.21 [0.34] | -3.54 | **<0.001** |
| Year (2011/2012) | -1.35 [0.33] | -4.09 | **<0.001** |
| Year (2012/2013) | -0.57 [0.33] | -1.72 | 0.086 |
| Year (2013/2014) | -1.61 [0.35] | -4.62 | **<0.001** |
| Year (2014/2015) | -1.15 [0.34] | -3.36 | **<0.001** |
| Year (2015/2016) | -0.86 [0.35] | -2.49 | 0.013 |
| Year (2016/2017) | -1.67 [0.35] | -4.72 | **<0.001** |
| Year (2017/2018) | -0.86 [0.36] | -2.39 | **0.017** |
| Year (2018/2019) | -1.19 [0.35] | -3.39 | **<0.001** |
| Year (2019/2020) | -0.47 [0.36] | -1.28 | 0.19 |
| Year (2020/2021) | -1.31 [0.37] | -3.57 | **<0.001** |
| Year (2021/2022) | -1.48 [0.37] | -4.02 | **<0.001** |
| Year (2022/2023) | -1.62 [0.36] | -4.55 | **<0.001** |
| Year (2023/2024) | -0.19 [0.38] | -0.51 | 0.61 |

**Table S5B. Pairwise habitat contrasts from the linear mixed effects model of the rate of morning weight gain (g/hr) presented in Table S5. For each season, the lower triangle of the matrix corresponds to the estimated difference in the marginal means [std error] and the upper triangle indicates the associated p-value. p values ≤ 0.05 are indicated in bold.**

| **May-Sep (DRY)** | Drie doring | Grassland | Red sand | White sand |
|---|---|---|---|---|
| Drie doring | -- | **0.039** | 0.82 | 0.37 |
| Grassland | 0.53 [0.20] | -- | **0.044** | **<0.001** |
| Red sand | 0.12 [0.14] | -0.41 [0.16] | -- | **<0.001** |
| White sand | -0.23 [0.14] | -0.76 [0.17] | -0.35 [0.08] | -- |
| **Oct-Apr (WET)** | Drie doring | Grassland | Red sand | White sand |
| Drie doring | -- | **0.023** | **0.009** | 0.080 |
| Grassland | 0.55 [0.19] | -- | 0.79 | **<0.001** |
| Red sand | 0.40 [0.13] | -0.14 [0.16] | -- | **<0.001** |
| White sand | -0.31 [0.13] | -0.86 [0.17] | -0.72 [0.08] | -- |

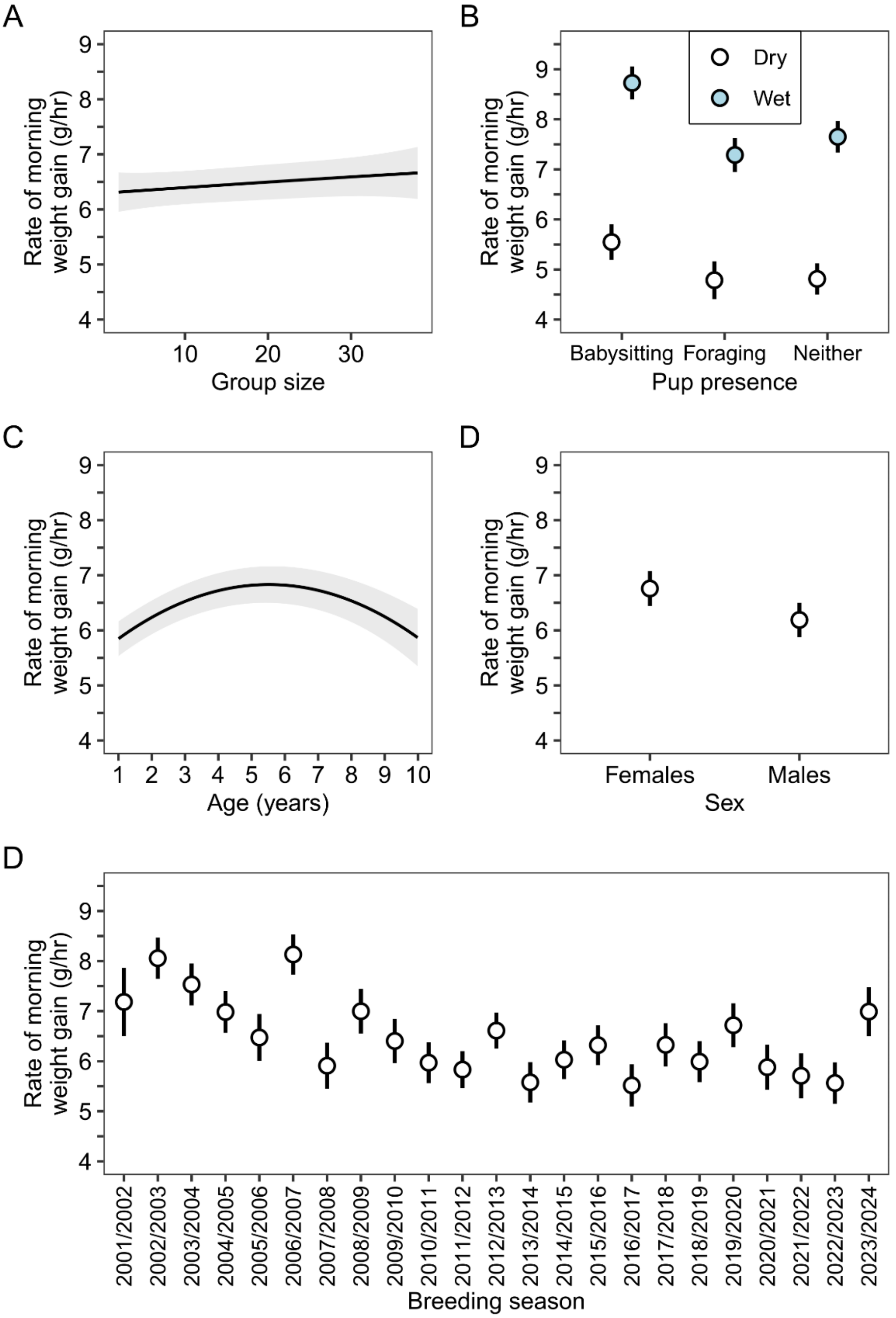


**Figure S5. Associations between the average rate of morning weight gain (g/hour) and (A) group size, (B) the presence of pups at the group, (C) individual age, (D) sex, and (E) the breeding season, as predicted by a linear mixed effects model. All panels display the predicted mean ± 95% confidence intervals.**

### *Group movement speeds*

**Table S6. Model output for the linear mixed effects model of group movement speeds (m/hour). The pairwise contrasts for period-specific habitat differences are presented in Table S5B. The intercept represents the reference habitat level of 'drie doring' in the dry season ('May-Sep') when a group was babysitting pups. We modelled the year as a fixed rather than a random effect so that we could estimate year-specific means directly (Figure S6C). The model included a random intercept of group identity (std dev = 25.06), date nested in group identity (std dev = 57.92), and a first order autocorrelation parameter (corAR1) was applied to the repeated observations within groups on a given morning (phi = 0.20). Residual std dev = 177.51. Continuous variables were standardised and centred for model fitting. p values ≤ 0.05 are indicated in bold.**

| Term | Estimate [Std Error] | t-value | p-value |
|---|---|---|---|
| Intercept | 234.92 [13.69] | - | - |
| Habitat (Grassland) | -33.87 [4.37] | -7.76 | **<0.001** |
| Habitat (Red sand) | -29.82 [2.78] | -10.73 | **<0.001** |
| Habitat (White sand) | 0.87 [2.66] | 0.33 | 0.74 |
| Season (Wet) | 21.05 [4.60] | 4.58 | **<0.001** |
| Habitat (Grassland) : Season (Wet) | 17.47 [5.58] | 3.13 | **0.002** |
| Habitat (Red sand) : Season (Wet) | 11.14 [3.57] | 3.12 | **0.002** |
| Habitat (White sand) : Season (Wet) | -1.15 [3.43] | -0.34 | 0.74 |
| Group size | 16.85 [1.27] | 13.30 | **<0.001** |
| Group size$^2$ | -3.65 [0.61] | -5.98 | **<0.001** |
| Time spent foraging | -35.16 [0.56] | -62.47 | **<0.001** |
| Time spent foraging$^2$ | 9.04 [0.54] | 16.58 | **<0.001** |
| Pup presence (neither) | -30.66 [3.38] | -9.08 | **<0.001** |
| Pup presence (foraging) | -44.79 [4.81] | -9.31 | **<0.001** |
| Pup presence (neither) : Season (Wet) | 14.14 [4.14] | 3.41 | **<0.001** |
| Pup presence (foraging) : Season (Wet) | -6.04 [5.75] | -1.05 | 0.29 |
| Year (2002/2003) | -19.27 [13.28] | -1.45 | 0.15 |
| Year (2003/2004) | -1.12 [13.21] | -0.08 | 0.93 |
| Year (2004/2005) | 5.74 [13.12] | 0.44 | 0.66 |
| Year (2005/2006) | 13.24 [13.63] | 0.97 | 0.33 |
| Year (2006/2007) | -39.93 [13.28] | -3.01 | **0.003** |
| Year (2007/2008) | -29.62 [13.51] | -2.19 | **0.028** |
| Year (2008/2009) | -20.08 [13.38] | -1.50 | 0.13 |
| Year (2009/2010) | -17.99 [13.81] | -1.30 | 0.19 |
| Year (2010/2011) | -11.89 [13.59] | -0.88 | 0.38 |
| Year (2011/2012) | -7.47 [13.35] | -0.56 | 0.58 |
| Year (2012/2013) | 39.19 [13.29] | 2.95 | **0.003** |
| Year (2013/2014) | 2.15 [13.30] | 0.16 | 0.87 |
| Year (2014/2015) | 26.7 [13.30] | 2.01 | **0.044** |
| Year (2015/2016) | 32.83 [13.33] | 2.46 | **0.014** |
| Year (2016/2017) | -11.31 [13.21] | -0.86 | 0.39 |
| Year (2017/2018) | -28.03 [13.39] | -2.09 | **0.036** |
| Year (2018/2019) | 6.67 [13.26] | 0.50 | 0.62 |
| Year (2019/2020) | 14.53 [13.41] | 1.08 | 0.28 |
| Year (2020/2021) | -20.39 [13.52] | -1.51 | 0.13 |
| Year (2021/2022) | -34.48 [13.60] | -2.54 | **0.011** |
| Year (2022/2023) | -67.63 [13.58] | -4.98 | **<0.001** |
| Year (2023/2024) | -1.80 [14.53] | -0.12 | 0.90 |

**Table S6B. Pairwise habitat contrasts from the the linear mixed effects model of group movement speeds during foraging (presented in Table S6). For each period, the lower triangle of the matrix corresponds to the estimated difference in the marginal means [std error] and the upper triangle indicates the associated p-value. p values ≤ 0.05 are indicated in bold.**

| **May-Sep (DRY)** | Drie doring | Grassland | Red sand | White sand |
|---|---|---|---|---|
| Drie doring | -- | **<0.001** | **<0.001** | 0.99 |
| Grassland | -33.87 [4.37] | -- | 0.69 | **<0.001** |
| Red sand | -29.82 [2.78] | 4.05 [3.71] | -- | **<0.001** |
| White sand | 0.87 [2.66] | 34.74 [4.03] | 30.69 [2.23] | -- |
| **Oct-Apr (WET)** | Drie doring | Grassland | Red sand | White sand |
| Drie doring | -- | **<0.001** | **<0.001** | 1.00 |
| Grassland | -16.40 [3.96] | -- | 0.91 | **<0.001** |
| Red sand | 18.68 [2.35] | -2.28 [3.42] | -- | **<0.001** |
| White sand | -0.28 [2.19] | 16.12 [3.72] | 18.40 [1.93] | -- |

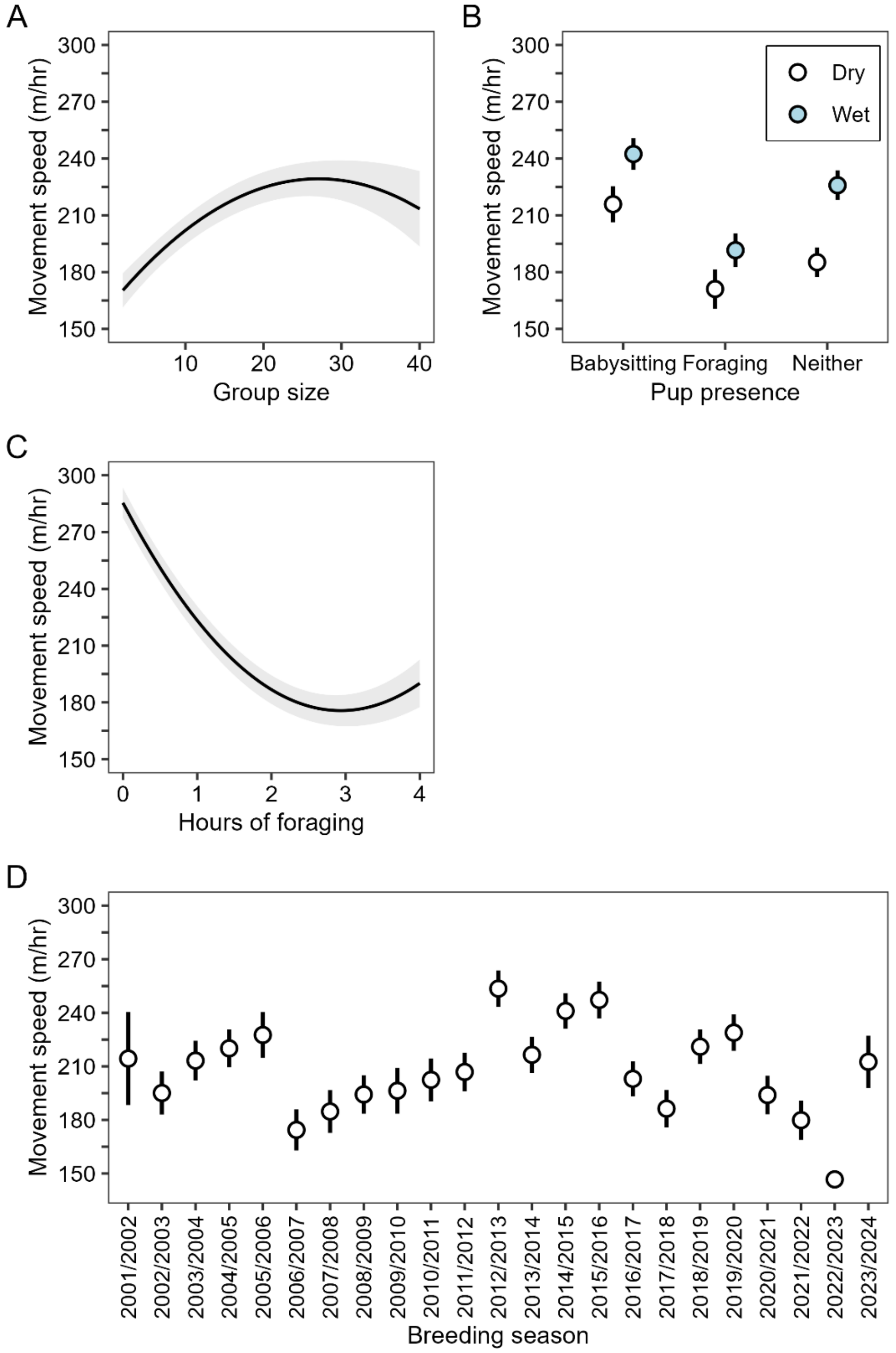


**Figure S6. Associations between the speed of group movements (m/hour) and (A) group size, (B) the presence of pups at the group, (C) the number of hours of morning foraging, and (D) the breeding season, as predicted by a linear mixed effects model. The presence of pups was either due to their permanent presence at burrow in early life, when they are babysat by one or more adult helpers, or later, once they begin foraging with the group. The number of hours foraging reflects the time since leaving the burrow in the morning. All panels display the predicted mean ± 95% confidence intervals. The model was fitted to individual movement steps from morning foraging tracks at approximately 15-minute intervals.**

### *Morning track lengths*

**Table S7. Model output for the linear mixed effects model fitted to morning track lengths (metres). The pairwise contrasts for period-specific habitat differences are presented in Table S7. The intercept represents the reference habitat level of 'drie doring' in the dry season ('May-Sep') when a group was babysitting pups. We modelled the year as a fixed rather than a random effect so that we could estimate year-specific means directly (Figure S7C). The model included a random intercept of group identity (std dev = 59.9), and an autocorrelation parameter (continuous autoregressive correlation of order 1, corCAR1) was applied to the repeated observations from groups over time (phi = 0.42). Residual std dev = 237.3. Continuous variables were standardised and centred for model fitting. p values ≤ 0.05 are indicated in bold.**

| Term | Estimate [Std Error] | t-value | p-value |
|---|---|---|---|
| Intercept | 629.9 [39.5] | - | - |
| Habitat (Grassland) | -24.6 [18.5] | -1.33 | 0.18 |
| Habitat (Red sand) | -35.9 [11.1] | -3.22 | **0.001** |
| Habitat (White sand) | -9.0 [9.7] | -0.93 | 0.35 |
| Season (Wet) | 84.8 [14.6] | 5.83 | **<0.001** |
| Habitat (Grassland) : Season (Wet) | -19.4 [22.6] | -0.86 | 0.39 |
| Habitat (Red sand) : Season (Wet) | -7.5 [13.9] | -0.53 | 0.59 |
| Habitat (White sand) : Season (Wet) | -13.6 [12.3] | -1.11 | 0.27 |
| Group size | 48.4 [3.7] | 13.0 | **<0.001** |
| Group size$^2$ | -10.8 [1.8] | -5.99 | **<0.001** |
| Total track duration | 85.4 [2.4] | 35.67 | **<0.001** |
| Pup presence (neither) | -82.8 [9.9] | -8.37 | **<0.001** |
| Pup presence (foraging) | -121.8 [14.2] | -8.58 | **<0.001** |
| Pup presence (neither) : Season (Wet) | 43.2 [12.2] | 3.54 | **<0.001** |
| Pup presence (foraging) : Season (Wet) | -12.2 [16.9] | -0.72 | 0.47 |
| Year (2002/2003) | -41.1 [38.9] | -1.06 | 0.29 |
| Year (2003/2004) | 6.2 [38.8] | 0.16 | 0.87 |
| Year (2004/2005) | 18.9 [38.5] | 0.49 | 0.62 |
| Year (2005/2006) | 34.4 [40.0] | 0.86 | 0.39 |
| Year (2006/2007) | -100.6 [39.0] | -2.58 | **0.010** |
| Year (2007/2008) | -51.8 [39.5] | -1.31 | 0.19 |
| Year (2008/2009) | -47.9 [39.2] | -1.22 | 0.22 |
| Year (2009/2010) | -43.6 [40.4] | -1.08 | 0.28 |
| Year (2010/2011) | -17.6 [39.7] | -0.44 | 0.66 |
| Year (2011/2012) | -7.0 [39.0] | -0.18 | 0.86 |
| Year (2012/2013) | 121.2 [38.9] | 3.12 | **0.002** |
| Year (2013/2014) | 24.1 [38.9] | 0.62 | 0.54 |
| Year (2014/2015) | 89.6 [38.9] | 2.30 | **0.021** |
| Year (2015/2016) | 113.0 [39.1] | 2.89 | **0.004** |
| Year (2016/2017) | -10.4 [38.8] | -0.27 | 0.79 |
| Year (2017/2018) | -55.7 [39.3] | -1.42 | 0.16 |
| Year (2018/2019) | 41.1 [38.9] | 1.06 | 0.29 |
| Year (2019/2020) | 58.9 [39.3] | 1.50 | 0.13 |
| Year (2020/2021) | -28.5 [39.5] | -0.72 | 0.47 |
| Year (2021/2022) | -60.9 [39.6] | -1.54 | 0.12 |
| Year (2022/2023) | -121.8 [39.7] | -3.07 | **0.002** |
| Year (2023/2024) | 21.4 [42.5] | 0.50 | 0.62 |

**Table S7B. Pairwise habitat contrasts from the linear mixed effects model fitted to morning tracks lengths (metres) presented in Table S7. For each season, the lower triangle of the matrix corresponds to the estimated difference in the marginal means [std error] and the upper triangle indicates the associated p-value. p values ≤ 0.05 are indicated in bold.**

| **May-Sep (DRY)** | Drie doring | Grassland | Red sand | White sand |
|---|---|---|---|---|
| Drie doring | -- | 0.54 | **0.007** | 0.79 |
| Grassland | -24.6 [18.5] | -- | 0.91 | 0.80 |
| Red sand | -35.9 [11.1] | -11.2 [17.0] | -- | **0.010** |
| White sand | -9.0 [9.7] | 15.6 [17.1] | 26.9 [8.7] | -- |

| **Oct-Apr (WET)** | Drie doring | Grassland | Red sand | White sand |
|---|---|---|---|---|
| Drie doring | -- | **0.044** | **<0.001** | **0.021** |
| Grassland | -44.1 [16.8] | -- | 1.00 | 0.54 |
| Red sand | -43.3 [9.2] | 0.7 [15.8] | -- | **0.035** |
| White sand | -22.6 [7.9] | 21.4 [16.1] | 20.7 [7.7] | -- |

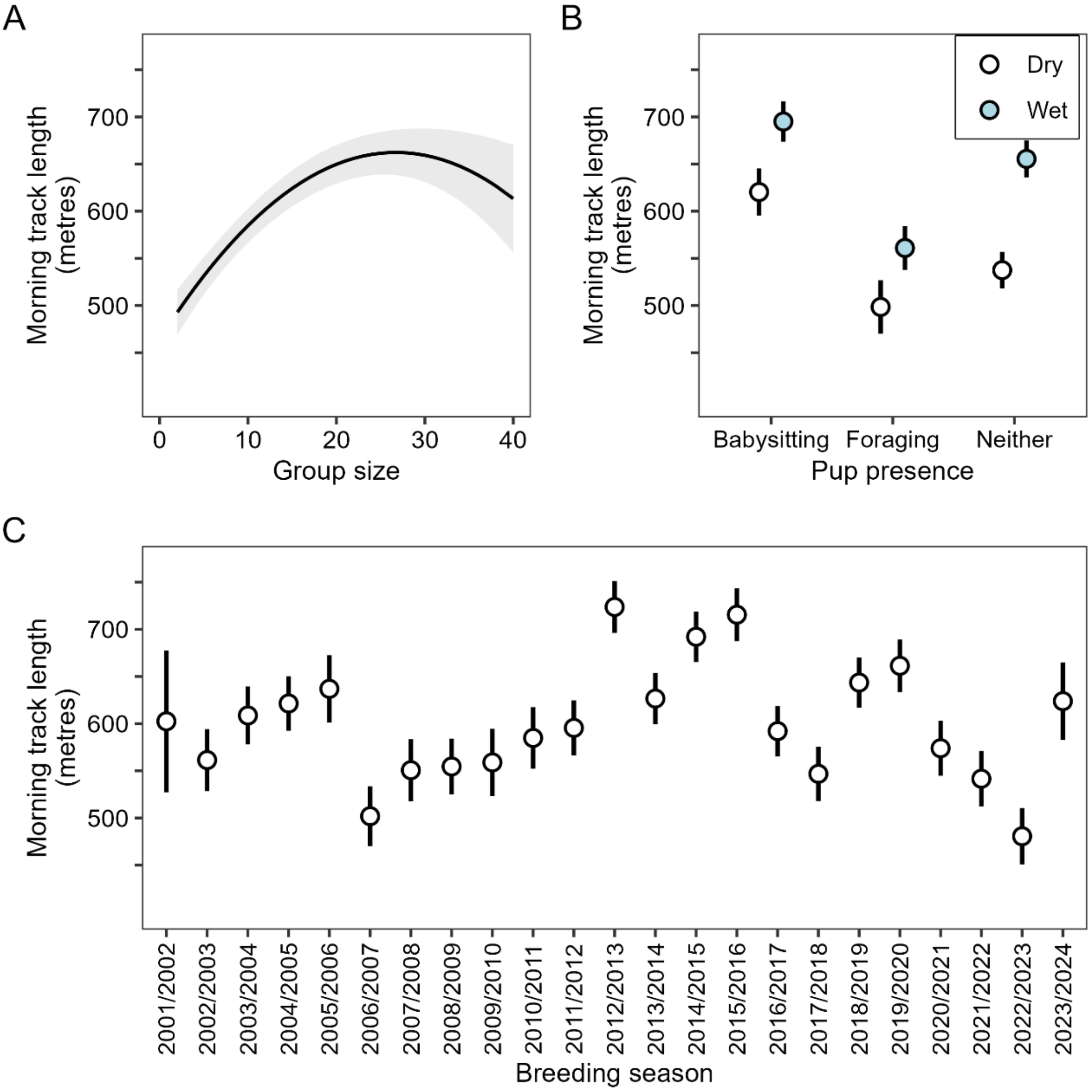


**Figure S7. Associations between the total morning track length and (metres) and (A) group size, (B) the presence of pups at the group, and (C) the breeding season, as predicted by a linear mixed effects model. All panels display the predicted mean ± 95% confidence intervals.**
